# MS2 RNA aptamer enhances prime editing in rice

**DOI:** 10.1101/2021.10.20.465209

**Authors:** Yiping Chai, Yuanyuan Jiang, Junya Wang, Dexin Qiao, Yu Zhang, Cuiping Xin, Yun Zhou, Xue-Chen Wang, Qi-Jun Chen

## Abstract

Prime editing is a universal and very promising precise genome editing technology. However, optimization of prime editor (PE) from different aspects remains vital for its use as a routine tool in plant basic research and crop molecular breeding. In this report, we tested MS2-based prime editor (MS2PE). We fused the M-MLV reverse transcriptase (RT) gene variant to the MS2 RNA binding protein gene, *MCP*, and allowed the MCP-RT fusion gene to co-express with the SpCas9 nickase gene, *SpCas9H840A*, and various engineered pegRNAs harboring MS2 RNA (MS2pegR). Compared with control PEs, MS2PEs significantly enhanced editing efficiency at four of six targets in rice protoplasts, and achieved 1.2∼10.1-fold increase in editing efficiency at five of six targets in transgenic rice lines. Furthermore, we tested total 22 different MS2pegR scaffolds, 3 RT variants or genes, 2 MCP variants, and various combinations of the Cas9 nickase, RT, and MCP modules. Our results demonstrated an alternative strategy for enhancing prime editing.

## Introduction

CRISPR/Cas systems that induced DNA double-strand breaks (DSBs) have been mainly used as a search-and-disrupt genome editing technology in plants (Zhan et al., 2021). DSB-based tools can also induce homology-directed repair (HDR) of host cells and thus can be used as a search-and-replace genome editing (Zhan et al., 2021). However, HDR-mediated gene targeting (GT) requires donor DNA, which in turn requires efficient delivery (Lu et al., 2020), synergetic processing (Barone et al., 2020), and in vivo amplifications or in vitro chemical modifications when possible (Cermak et al., 2015; Lu et al., 2020). In addition, as non-homologous end joining (NHEJ) is the preferred DSB repair mechanism in somatic plant cells, inherent low efficiency of HDR has been the main obstacle to practical applications of GT in crops (Zhan et al., 2021). These aspects prevent GT from broad applications by plant researchers although major advances have been made (Barone et al., 2020; Lu et al., 2020).

CRISPR/Cas-derived base editing without requiring DSBs or donor DNA templates is a search-and-convert editing technology for base conversions (Koblan et al., 2018; Jin et al., 2020; Richter et al., 2020). However, although two types of base editors for inducing all 4 types of base transition mutations and a type of base editors for inducing C to G transversion in mammalian cells have been developed, base editors for inducing the rest 6 types of base transversion mutations remain to be developed (Anzalone et al., 2019; Kurt et al., 2021; Zhao et al., 2021). In addition, base editing has a much stricter requirement for target selection due to the restriction of editing windows and has the potential to induce off-target mutations in editing windows harboring multiple editable bases. These aspects restrict broad applications of base editors.

Like base editing, CRISPR/Cas-derived prime editing requires no DSBs or donor DNA for precise genome modifications. However, unlike base editing, which is mainly used for inducing base conversions, prime editing is a universal search-and-replace genome modification technology (Anzalone et al., 2019). Prime editors (PEs) can induce controllable rather than random indel mutations and unlimited base conversions, and thus has a potency to substitute the HDR-based genome editors to a large extent and base editors in a complete way. In plants, the main obstacle to broad applications of prime editing is the overall low editing efficiency of PEs (Jiang et al., 2020; Lin et al., 2021). Thus, optimization of PEs for higher editing efficiency from different aspects is vital for their acceptance as routine tools by plant researchers. In this report, we developed MS2 RNA aptamer-based prime editing in rice and demonstrated that the MS2-based strategy greatly improved prime-editing efficiency.

## Results and Discussion

### MS2-based prime editors enhance prime editing in rice protoplasts and transgenic lines

The MS2 RNA aptamer and its binding protein MCP in combination with CRISPR/Cas9 system have been used for efficient gene activation and base editing (Konermann et al., 2015; Zalatan et al., 2015; Hess et al., 2016; Li et al., 2020), however, it remains unknown whether the MS2 system can be used for efficient prime editing. To test MS2-based prime editors (MS2PEs), as an initial step we tested five engineered pegRNA scaffolds, named MS2pegRs. These five MS2pegRs harbor two copies of MS2 RNA aptamers, MS2 and/or f6, which are located at different regions of the pegRNA scaffold (Figure 1a, b; Figure S1). We fused the M-MLV reverse transcriptase (RT) gene variant to *MCP* and allowed the *MCP-RT* fusion gene to co-express with a Cas9 nickase gene, *SpCas9H840A*, and MS2pegR genes (Figure 1a, c). We tested two MCP variants, MCP1 and MCP2 (Supplemental information), and the five MS2pegR scaffolds at six rice genomic targets.

**Figure 1.**
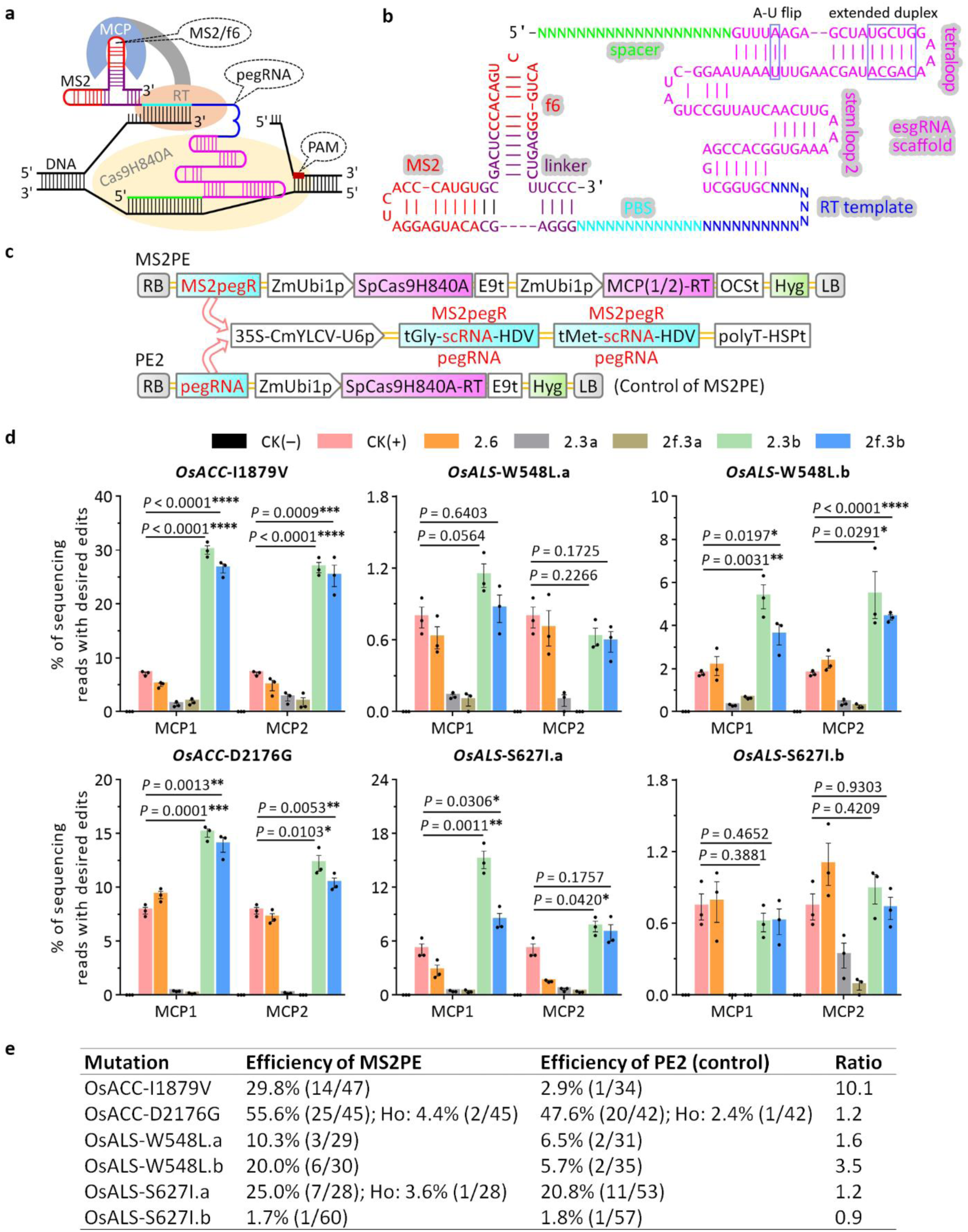
MS2 RNA aptamer enhances prime editing in rice. **a**, Schematic diagram of MS2-based prime editing. **b**, Secondary structure of a representative engineered pegRNA (pegR2f.3b) harboring MS2 and f6 RNA aptamers. esgRNA, enhanced sgRNA with modifications indicated. Besides 3’-end, the other two regions, tetraloop and stem loop 2 of the sgRNA scaffold used for inserting MS2 and f6 hairpins are indicated. PBS, primer binding site. **c**, Structure of PEs. MCP1/2, two MCP variants. scRNA, CRISPR scaffold RNA. tGly/Met, tRNA(Gly/Met). E9t, OCSt, and HSPt, terminators from pea, Agrobacterium, and Arabidopsis, respectively. **d**, Efficiencies of MS2 prime editors with different MS2pegRNA scaffolds and MCP variants in rice protoplasts. Each MS2 prime editor harbors one of five different MS2pegRNA scaffolds (pegRNA2.6, 2.3a, 2f.3a, 2.3b, and 2f.3b) and one of two MCP variants (MCP1 and 2). W548L.a/b or S627I.a/b represent 2 completely different MS2pegRNAs inducing the same mutations. Untreated protoplast samples served as negative controls named CK(–) and protoplast samples treated with original prime editors served as positive controls named CK(+). Efficiency (mean ± s.e.m.) was calculated from three independent experiments (n = 3). *P* values were obtained by using the two-tailed Student’s *t*-test, comparing MS2PEs with positive controls. \**P* < 0.05, \*\**P* < 0.01, \*\*\**P* < 0.001, \*\*\*\**P* < 0.0001. **e**, Efficiencies of mutations induced by PEs in rice transgenic lines. The ratios of T0 mutants to total number of T0 transgenic plants are indicated in parentheses. Ho, efficiency of homozygous mutations when available.

We first tested editing efficiency of MS2PEs in rice protoplasts. Compared with control PEs, out of the five MS2pegR scaffolds, pegR2.3b and pegR2f.3b (Figure S1) significantly enhanced editing efficiency at four of six targets, and displayed similar editing efficiency at the rest two targets (Figure 1d). In general, there was no significant difference between pegR2.3b and pegR2f.3b. There was also no significant difference between MCP1 and MCP2 (Figure 1d). We further tested editing efficiency of MS2PEs in rice transgenic lines (Figure 1e). The results indicated that MS2PEs, each of which harbors two similar MS2pegRs, pegR2.3b and pegR2f.3b, for enhancing expression, increased editing efficiencies by a factor of 1.2∼ 10.1 times at five of six targets compared with control PEs, each of which harbors two same pegRNAs for enhancing expression (Figure 1c, e). These results demonstrated that MS2PEs greatly enhanced prime editing in rice.

In protoplasts, the most efficient three MS2PEs were those inducing mutations in OsACC-I1879V, OsACC-D2176G, and OsALS-S627I.a, whereas in transgenic lines, the most efficient three MS2PEs were those inducing mutations in OsACC-D2176G, OsACC-I1879V, and OsALS-S627I.a. This inconsistency may reflect on differences of edited cell types, i.e., protoplast cells and callus cells. In theory, editing processes last for much longer time in callus cells than in protoplast cells and promoters may have different activity in these two types of cells.

### Comparisons of various MS2PEs with different MS2pegR scaffolds and RT modules

To attempt to optimize MS2PEs, we tested more MS2PEs with different MS2pegR scaffolds and RT modules. To more extensively evaluate effects of MS2pegRNA scaffolds on editing efficiency, we designed additional 17 MS2pegR scaffolds (Figure 2a; Figure S2-S6). The results indicated that only pegR1.3b and pegR1f.3b achieved editing efficiency similar to pegR2.3b or pegR2f.3b (Figure 2a, b). These results suggested that regions which MS2 and f6 located are decisive factors for editing efficiency whereas copy number of MS2 and f6 is not an important factor (Figure 2a; Figure S2-S6).

**Figure 2.**
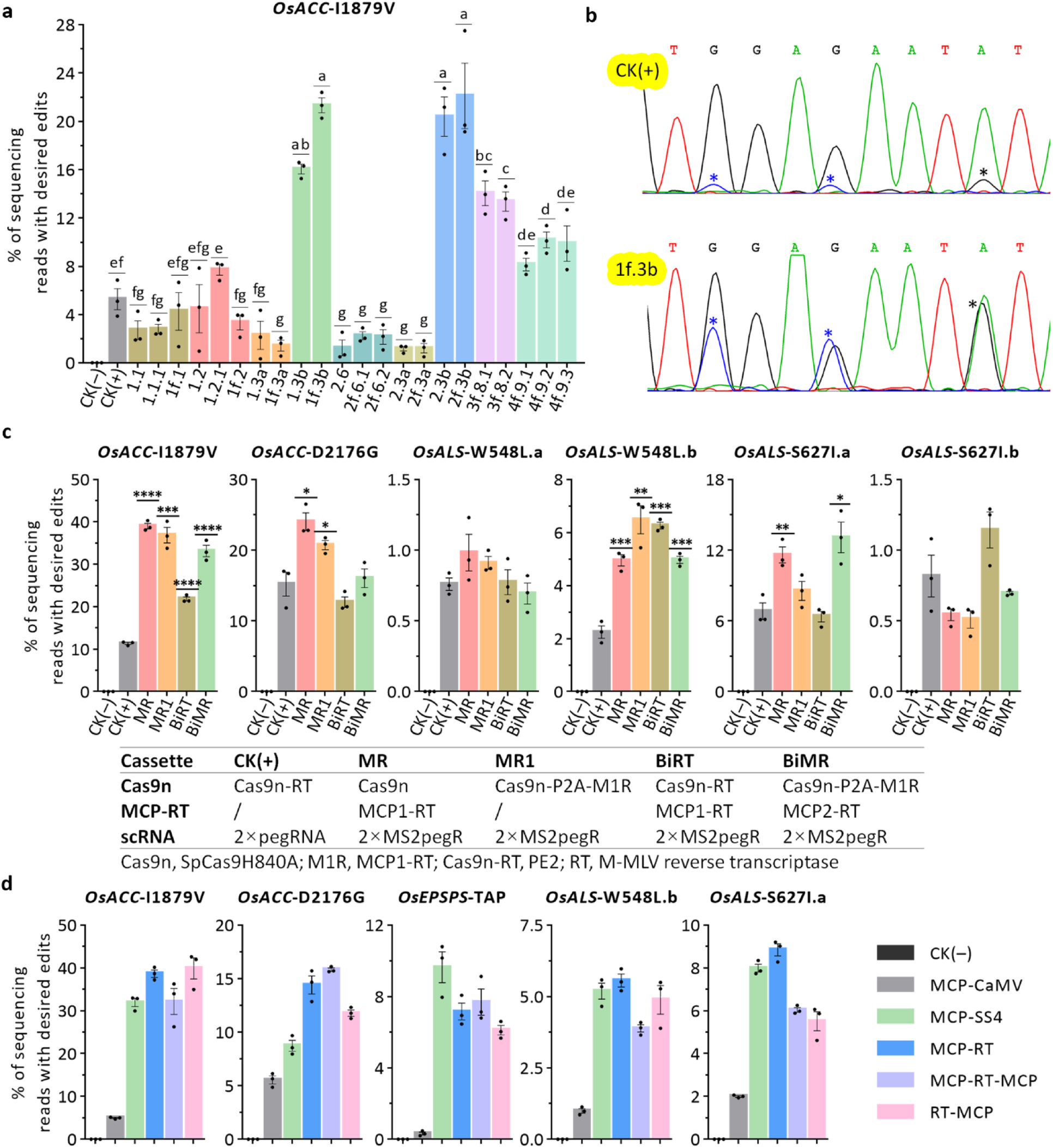
Comparisons of various MS2PEs in rice protoplasts. **a**, Comparisons of various MS2PEs harboring different MS2pegRNA scaffolds. Each MS2 prime editor harbors one of 22 different MS2pegRNA scaffolds. Efficiency (mean ± s.e.m.) was calculated from three independent experiments (n = 3). The absence of the same letter indicates a significant difference, *P* values were obtained by using the two-tailed Student’s *t*-test (*P* < 0.05). **b**, High-frequency mutations induced by PEs in rice protoplasts could be detected from sequencing chromatograms of the PCR fragments amplified from protoplast DNA samples. Two representatives are shown and an asterisk indicates a mutation induced by PEs. **c**, Comparisons of MS2PEs with different RT-containing modules or vector structures. Each MS2PE harbors the same MS2pegRNA scaffold (pegRNA2f.3b) and a Cas9n cassette with or without the MCP-RT cassette. The cassettes in the five PEs are indicated. Untreated protoplast samples served as negative controls named CK(–) and protoplast samples treated with original PEs served as positive controls named CK(+). Efficiency (mean ± s.e.m.) was calculated from three independent experiments (n = 3). The *P* values were obtained by using the two-tailed Student’s *t*-test, comparing MS2PEs with positive controls. \**P* < 0.05, \*\**P* < 0.01, \*\*\**P* < 0.001, \*\*\*\**P* < 0.0001. **d**, Comparisons of various combinations of MCP and RT by using a dual vector-based strategy for protoplast transfection. In the dual vector system, one vector harbors the Cas9n cassette and two same MS2pegRNAs whereas another vector harbors a fusion gene of MCP and RT.

Comparison of MS2PEs with different structures found that MS2PEs with a single cassette harboring the fusion gene of Cas9n and MCP-RT, had similar efficiency to those with two cassettes driving the expression of Cas9n and MCP-RT separately (Figure 2c). Comparison of MS2PEs with additional RT modules found that additional RT modules had no contributions to improving MS2PEs (Figure 2c). Comparison of MS2PEs with different fusion ways of RT and MCP found that MCP-RT fusion gene showed similar overall editing efficiency to MCP-RT-MCP and RT-MCP fusion genes (Figure 2d). These results suggested that MCP positions relative to RT have no effect on MS2PE activity, and additional MCP have no contributions to improving MS2PEs. Comparison of MS2PEs with different RT genes or variants found that the SuperScript4 (SS4) M-MLV RT variant (Thermo Fisher Scientific) harboring 20 mutated amino acids showed similar editing efficiency to the RT variant from PE2, and that CaMV-RT showed much lower editing efficiency than the M-MLV RT variants (Figure 2d). We anticipate that further evolution of SS4 RT variant for temperature-tolerant versions like ttLbCas12a (Schindele and Puchta, 2020) will facilitate the improvement of MS2PEs.

In conclusion, we showed that MS2-based PEs greatly enhance prime editing efficiency in rice and demonstrated the decisive factors for high efficiency of MS2PEs. We presented an alternative strategy for enhancing prime editing and provided an alternative platform for further directed evolution of RT variants or genes. This strategy also facilitates the development of additional PEs based on other orthogonal CRISPR/Cas systems.

## Methods

All primers used in this study are listed in Table S1, and the sequences of the MCP-RT and pegRNA expression cassettes are listed in Supplemental information. Vectors described in this study together with their annotated sequences are available from Addgene and/or MolecularCloud (GenScript).

### Vector construction

We amplified RT from pUC57-PE2 with primers RT-NXF/-SaR, purified the PCR products, digested them with *Nco*I and *Sac*I, and allowed them to ligate with *Nco*I and *Sac*I-digested pR1R4-UBQ1-U6A, which resulted in the generation of pR1R4-RT. We replaced the *Asc*I-*Eco*RI fragment of pL2R4-U6-mCh with the *Asc*I-*Eco*RI fragment of pR1R4-RT, resulting in the generation of pL2R4-RT. We replaced the *Nco*I-*Bst*XI fragment of pL2R4-RT with a synthetic *MCP1* (Zalatan et al., 2015) or *MCP2* (Konermann et al., 2015) fragment digested with *Nco*I and *Bst*XI, resulting in the generation of pL2R4-MCP-RT or pL2R4-MCP2-RT, respectively. We replaced the *Pme*I-*Spe*I fragment of pG3R23-PE2-U3A.2 (Jiang et al., 2020) with an insert prepared by annealing two oligos, oPiScH-F/-R, resulting in the generation of pG3R23-Ubi. We replaced the *Hin*dIII-*Xba*I fragment of pL2R4-MCP-RT or pL2R4-MCP2-RT with the *Hin*dIII-*Xba*I fragment of pG3R23-Ubi, resulting in the generation of <u>pL2R4-ZmUbi-MR</u> or <u>pL2R4-ZmUbi-M2R</u>, respectively. We removed the *Asc*I-*Eco*RI fragment of pL4L3-U3H.2 (Jiang et al., 2020), resulting in the generation of <u>pL4L3-Hyg2</u>. We replaced the *Xba*I-*Sac*I fragment of PE2 on pG3R23-PE2-35C (Jiang et al., 2020) with the *Xba*I-*Sac*I fragment of pUC57-zCas9H840A (Xing et al., 2014), resulting in the generation of <u>pG3R23-840-35C</u>.

We used MultiSite Gateway technology to assemble MS2pegRNA-cloning MS2 PE vectors. We mixed vectors including pG3R23-840-35C, pL2R4-ZmUbi-MR, and pL4L3-Hyg2, and reagents of MultiSite Gateway kit (Invitrogen) to assemble <u>pG3H-MR-35C</u>. We replaced pL2R4-ZmUbi-MR of the above three vectors with pL2R4-ZmUbi-M2R to assemble <u>pG3H-M2R-35C</u>. To generate final MS2-based prime editors, we replaced the *Bsa*I fragments of the above MS2pegRNA-cloning vectors with *Bsa*I-digested synthetic fragments harboring two MS2pegRs, resulting in the generation of MR and M2R vectors. To generate control prime editors, we replaced the *Bsa*I fragments of pG3H-PE2-35C (Jiang et al., 2020) with *Bsa*I-digested synthetic fragments harboring two same pegRNAs, resulting in the generation of CK(+) vectors.

We replaced the *Hin*dIII-*Sal*I fragment of pL2R4-MCP-RT with a synthetic fragment digested with *Hin*dIII and *Sal*I, resulting in the generation of pL2R4-P2A-MR. We replaced the *Mlu*I-*Sac*I fragment of p35C-ALS-S1 (Jiang et al., 2020) with a synthetic fragment, resulting in the generation of pG3H-840P1. We replaced the *Bsa*I-*Sac*I fragment of pG3H-840P1 with the *Bsa*I-*Sac*I fragment of pL2R4-P2A-MR, resulting in the generation of pG3H-840MR1. We replaced the *Hin*dIII-*Spe*I fragment of pG3H-840MR1 with the *Hin*dIII-*Spe*I fragment of pG3R23-PE2-35C, resulting in the generation of <u>pG3H-MR1-35C</u>. We replaced the *Eco*RI-*Sac*II fragment of pG3H-MR1-35C with the *Eco*RI-*Sac*II fragment of pG3R23-PE2-35C, resulting in the generation of <u>pG3R23-MR1-35C</u>. We mixed vectors including pG3R23-PE2-35C, pL2R4-ZmUbi-MR, and pL4L3-Hyg2, and reagents of MultiSite Gateway kit (Invitrogen) to assemble <u>pG3H-BiRT-35C</u>. We mixed vectors including pG3R23-MR1-35C, pL2R4-ZmUbi-M2R, and pL4L3-Hyg2, and reagents of MultiSite Gateway kit (Invitrogen) to assemble <u>pG3H-BiMR-35C</u>. To generate final MS2-based prime editors, we replaced the *Bsa*I fragments of pG3H-MR1-35C, pG3H-BiRT-35C, or pG3H-BiMR-35C with *Bsa*I-digested synthetic fragments harboring two same MS2pegRs, resulting in the generation of MR1, BiRT, or BiMR vectors.

To use dual vector-based strategy for comparisons of various MR modules in protoplasts, we replaced the *Eco*RI-*Sac*II fragment of pG3R23-840-35C with an insert prepared by annealing two oligos, resulting in the generation of <u>pG3-840-35C</u>. We replaced the *Xho*I-*Sac*I fragment of pL2R4-MCP-RT with a synthetic fragment digested with *Xho*I and *Sac*I, resulting in the generation of <u>pL2R4-MRM</u>. We replaced the *Xba*I-*Bst*XI fragment of pL2R4-MRM with a synthetic fragment digested with *Xba*I and *Bst*XI, resulting in the generation of <u>pL2R4-RM</u>. We replaced the *Asc*I-*Xba*I fragment of pL2R4-MRM or pL2R4-RM with ZmUbi1 promoter digested with *Asc*I and *Xba*I, resulting in the generation of <u>pL2R4-Ubi-MRM</u> or <u>pL2R4-Ubi-RM</u>. We replaced the *Xba*I-*Sac*I fragment of pL2R4-Ubi-MRM with a synthetic MCP-CaMV_RT or MCP-SS4_RT digested with *Xba*I and *Sac*I, resulting in the generation of <u>pL2R4-Ubi-CaMV</u> (Lin et al., 2020) or <u>pL2R4-Ubi-SS4</u>. To clone MS2pegRs, we replaced the *Bsa*I fragment of pG3-840-35C with *Bsa*I-digested synthetic fragments harboring two same MS2pegRs. For dual vector-based protoplast transfection, we used a pG3-840-35C-derived vector harboring Ubi-Cas9n and MS2pegRs, and one of the five vectors including pL2R4-ZmUbi-MR (MCP-RT), pL2R4-Ubi-MRM (MCP-RT-MCP), pL2R4-Ubi-RM (RT-MCP), pL2R4-Ubi-CaMV (MCP-CaMV_RT), and pL2R4-Ubi-SS4 (MCP-SS4_RT) to perform rice protoplast transfection.

### Rice protoplast transfection and analysis of prime editing

We used the Japonica rice (*Oryza sativa*) variety Zhonghua11 to prepare protoplasts. We transferred the plasmids into protoplasts by PEG-mediated methods and incubated the transfected protoplasts at 26 °C for 48 hours. To analyze efficiency of prime editing, we extracted the genomic DNA and used the primers listed in Table S1 to amplify target fragments for deep sequencing. For each target site, sequencing fragment was repeated three times using genomic DNA extracted from three independent protoplast samples.

### Rice transformation and analysis of prime editing

We separately transformed the pGreen3 binary vectors into the engineered Agrobacterium strain LBA4404/pVS1-VIR2 to generate strains harboring the ternary vector system (Zhang et al., 2019). We used these strains to transform callus cells of Zhonghua11 separately. To analyze the mutations induced by prime editing, we amplified fragments spanning the target sites in genes from genomic DNA of the transgenic lines using PCR with primers listed in Table S1. We then submitted the purified PCR products to direct sequencing with primers listed in Table S1.

## Author contributions

Q.J.C., Y.Zhou, and X.C.W. designed the research. Y.C., Y.J., J.W., D.Q., Y.Z., and C.X. performed the experiments. Q.J.C., Y.C., Y.J., Y.Zhou, and X.C.W. wrote the manuscript. Q.J.C. supervised the project.

## Acknowledgements

This work was supported by grants from the National Natural Science Foundation of China (grant nos. 31872678, U19A2022, and 31670371), National Transgenic Research Project (grant nos. 2019ZX08010003-001-011 and 2016ZX08009002), and the National Crop Breeding Fund (grant no. 2016YFD0101804).

