## Supplemental information for "MS2 RNA aptamer enhances prime editing in rice"

### Table of contents

Supplementary Figure S1 ... 4
Supplementary Figure S2 ... 5
Supplementary Figure S3 ... 6
Supplementary Figure S4 ... 7
Supplementary Figure S5 ... 8
Supplementary Figure S6 ... 9
Supplementary Table S1. Sequences of primers ... 10
Supplemental sequence 1. MCP1-RT ... 11
Supplemental sequence 2. MCP2-RT ... 11
Supplemental sequence 3. MCP-RT-MCP ... 11
Supplemental sequence 4. RT-MCP ... 12
Supplemental sequence 5. SpCas9H840A-P2A-MR ... 12
Supplemental sequence 6. MCP-CaMV_RT ... 12
Supplemental sequence 7. MCP-SS4_RT ... 13
Supplemental sequence 8. Synthetic 2xMS2pegRs or 2xpegRNAs ... 13
Supplemental sequence 9. Cassette expressing two MS2pegRs or pegRNAs ... 13
Supplemental sequence 10. MS2pegR scaffolds ... 14
- >pegRNA ... 14
- >pegRNA2.6 ... 14
- >pegRNA2.3a ... 14
- >pegRNA2f.3a ... 14
- >pegRNA2.3b ... 14
- >pegRNA2f.3b ... 14
- >pegRNA1.1 ... 14
- >pegRNA1.2 ... 14
- >pegRNA1.1.1 ... 14
- >pegRNA1.2.1 ... 14
- >pegRNA1f.1 ... 14
- >pegRNA1f.2 ... 14
- >pegRNA1.3a ... 14
- >pegRNA1f.3a ... 15
- >pegRNA1.3b ... 15
- >pegRNA1f.3b ... 15
- >pegRNA2f.6.1 ... 15
- >pegRNA2f.6.2 ... 15
- >pegRNA3f.8.1 ... 15
- >pegRNA3f.8.2 ... 15
- >pegRNA4f.9.1 ... 15
- >pegRNA4f.9.2 ... 15
- >pegRNA4f.9.3 ... 15

Supplemental sequence 11. Spacer, rtT, and PBS of MS2pegRs ... 16
- >OsACC-I1879V ... 16

#### Supplementary Figure S1

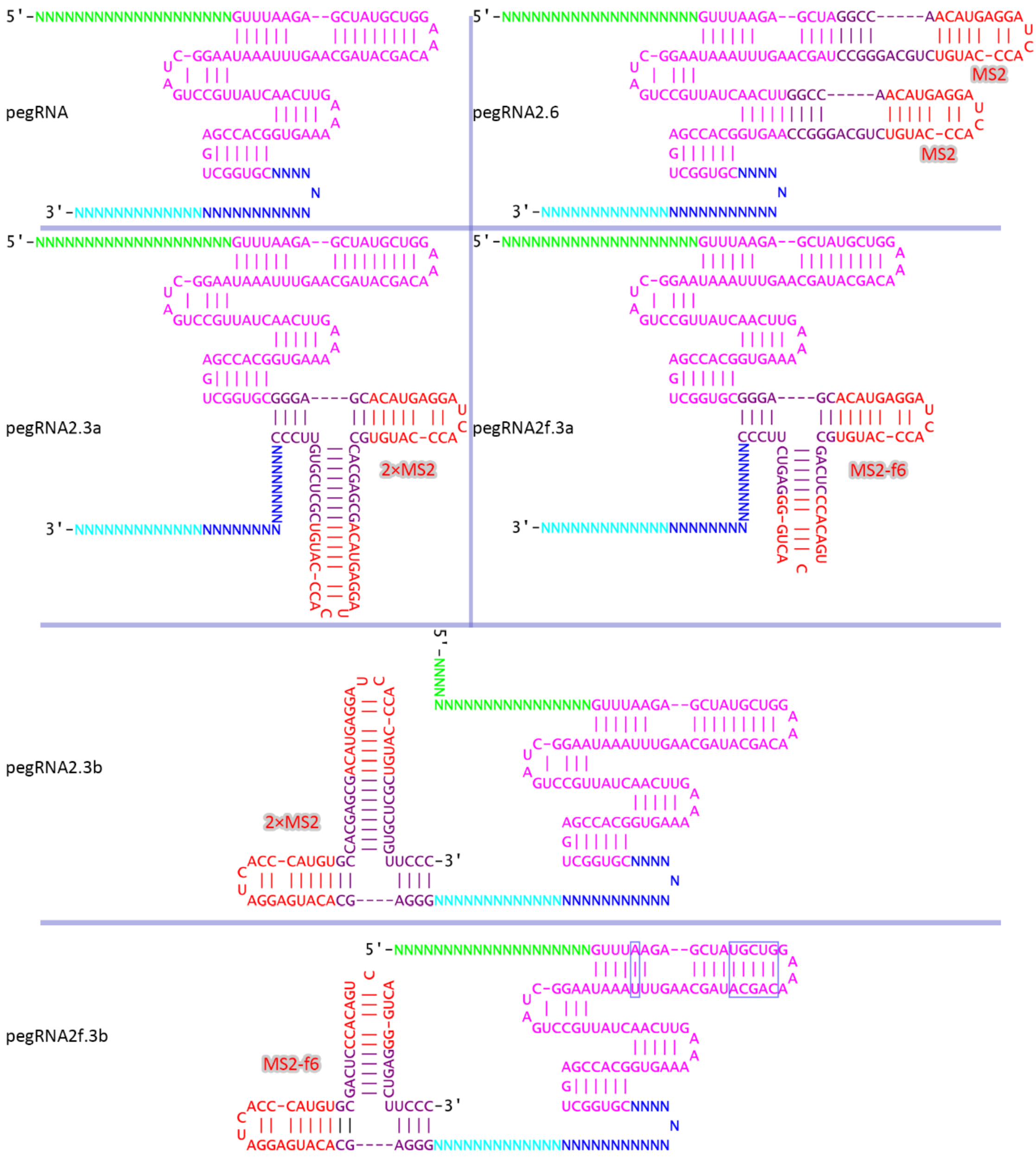

**Supplementary Figure S1. Secondary structures of 5 engineered pegRNA scaffolds including pegRNA2.6, pegRNA2.3a/b, and pegRNA2f.3a/b.**

For nomenclature, the first digit 2 represents two MS2 and f6 hairpins and “f” following the first digit indicates existence of f6. The second digit 3 represents 3’-end of sgRNA or pegRNA and “a” or “b” following the digit 3 represents 3’-end of sgRNA or pegRNA, respectively. The second digit 6 represents the two regions of tetraloop and stem loop 2.

#### Supplementary Figure S2

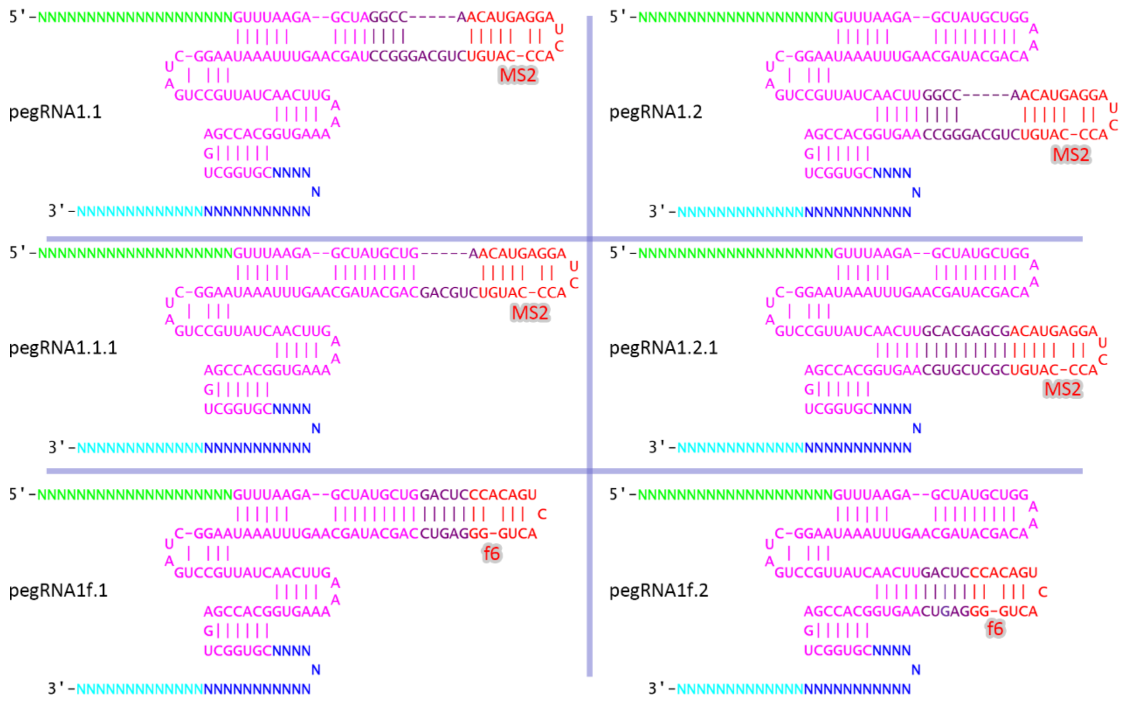

**Supplementary Figure S2. Secondary structures of 6 engineered pegRNA scaffolds including pegRNA1.1, pegRNA1.2, pegRNA1.1.1, pegRNA1.2.1, pegRNA1f.1, and pegRNA1f.2.**

For nomenclature, the first digit 1 represents one MS2 or f6 hairpin and “f” following the first digit indicates existence of f6. The second digits 1 and 2 represent tetraloop and stem loop 2 of sgRNA scaffold, respectively. The third digit 1 represents a different linker between sgRNA scaffold and MS2 or f6.

#### Supplementary Figure S3

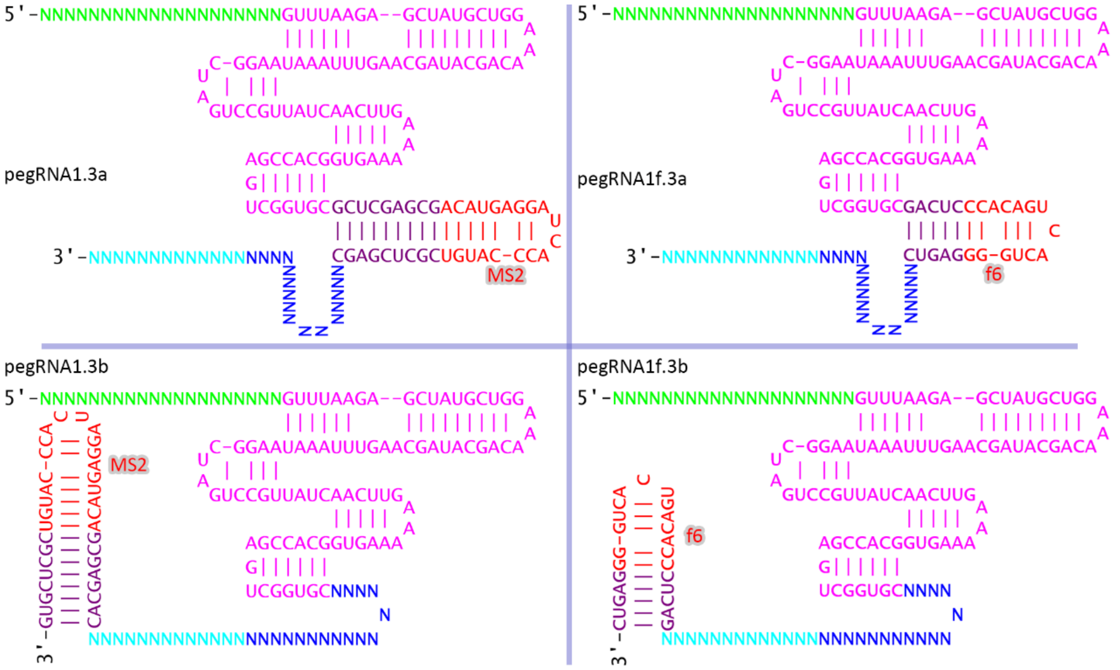

**Supplementary Figure S3. Secondary structures of 4 engineered pegRNA scaffolds including pegRNA1.3a/b and pegRNA1f.3a/b.**

For nomenclature, the first digit 1 represents one MS2 or f6 hairpin and "f" following the first digit indicates existence of f6. The second digit 3 represents 3'-end of sgRNA or pegRNA; "a" or "b" following the digit 3 represents 3'-end of sgRNA or pegRNA, respectively.

#### Supplementary Figure S4

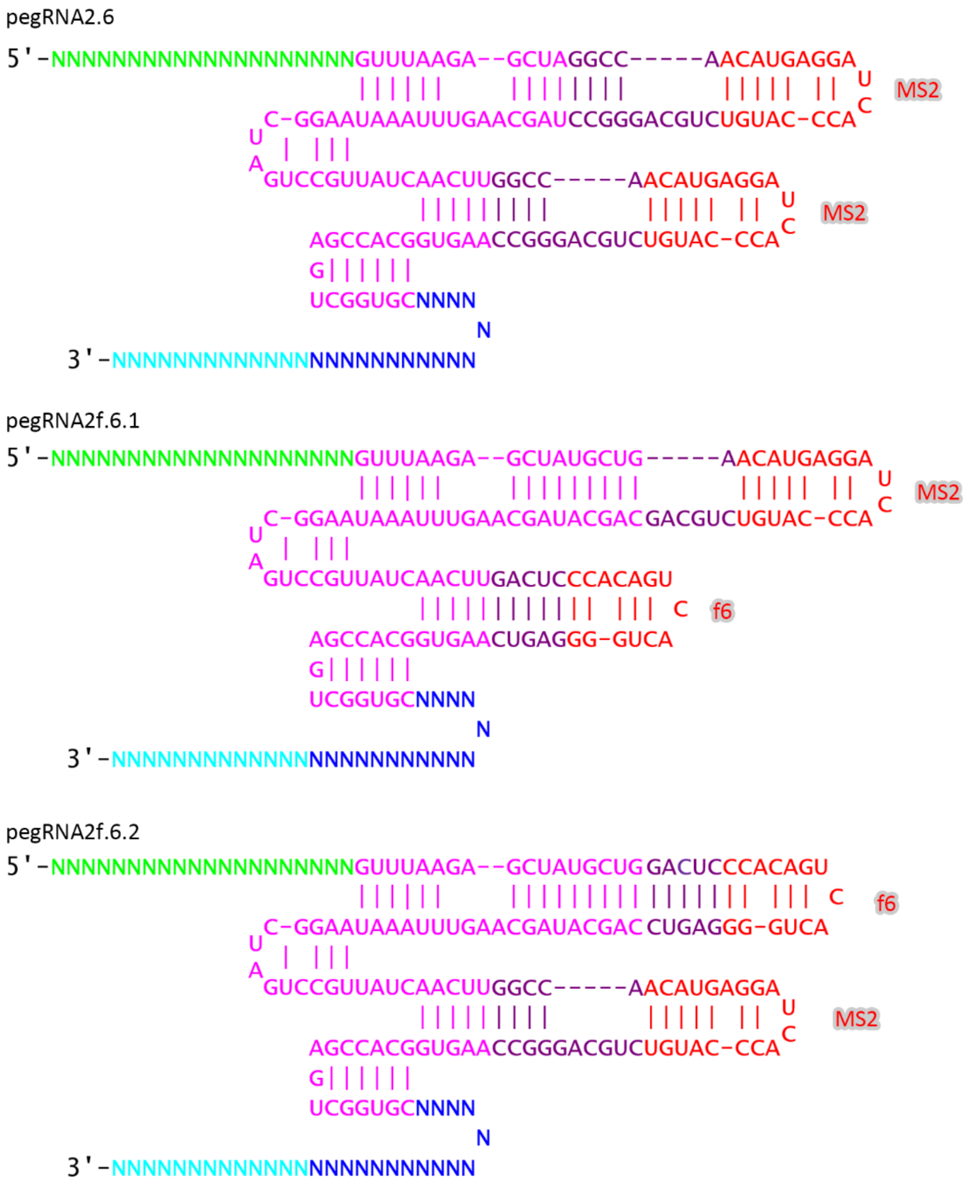

**Supplementary Figure S4. Secondary structures of 2 engineered pegRNA scaffolds including pegRNA2f.6.1/2.** For nomenclature, the first digit 2 represents two MS2 and f6 hairpins and "f" following the first digit indicates existence of f6. The second digit 6 represents the two regions of tetraloop and stem loop 2 of sgRNA scaffold. The third digits represent other differences.

#### Supplementary Figure S5

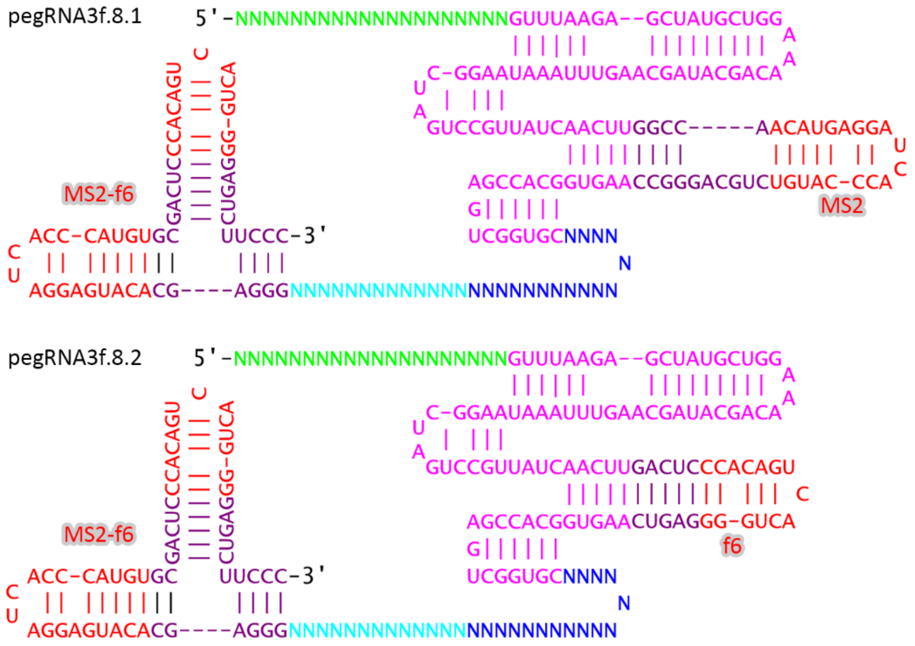

**Supplementary Figure S5. Secondary structures of 2 engineered pegRNA scaffolds including pegRNA3f.8.1/2.** For nomenclature, the first digit 3 represents three MS2 and f6 hairpins and “f” following the first digit indicates existence of f6. The second digit 8 represents the two regions of stem loop 2 and 3’-end of pegRNA scaffold. The third digits represent other differences.

### Supplementary Figure S6

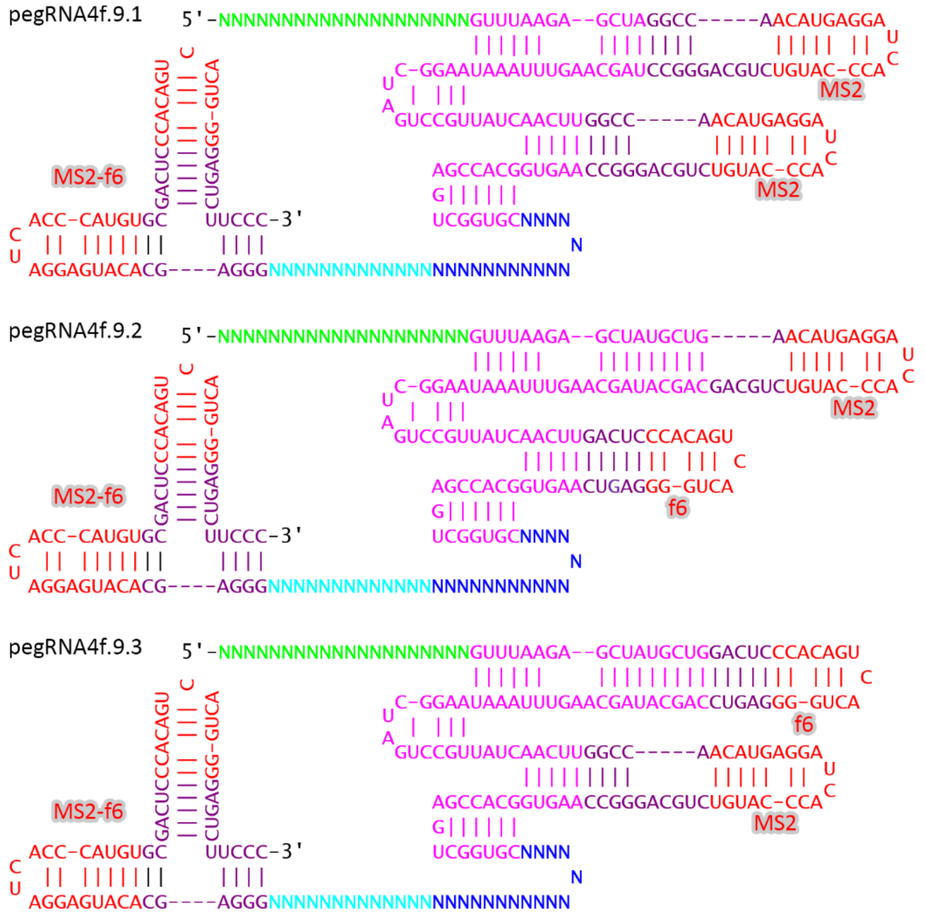

**Supplementary Figure S6. Secondary structures of 3 engineered pegRNA scaffolds including pegRNA4f.9.1/2/3.** For nomenclature, the first digit 4 represents four MS2 and f6 hairpins and "f" following the first digit indicates existence of f6. The second digit 9 represents the three regions of tetraloop, stem loop 2, and 3'-end. The third digits represent other differences.

### Supplementary Table S1. Sequences of primers

**Supplementary Table S1.** Sequences of primers

| Sequence name | Sequence | Purpose |
|---|---|---|
| RT-NXF | ATATTTCCATGGTCTAGATGACCCTGAACATTGAGGAC | Vector construction |
| RT-SaR | ATTTATTGAGCTCACACCTTCCTCTTCTTC | |
| oPiScH-F | AAACATTACCCTGTTATCCCTAAGCTTG | |
| oPiScH-R | CTAGCAAGCTTAGGGATAACAGGGTAATGTTT | |
| ALSWS-IDF | CTAACCCAGGTGTCACAGTTGTT | Analysis of the W548L and S627I mutations by Sanger sequencing |
| ALSWS-IDR | AGAGCACATACAAACATCATAGGC | |
| I1879V-IDF | AGTGAAATCTTGCTTCCGTGTTG | Analysis of the I1879V mutation by Sanger sequencing |
| I1879V-IDR | AATGTAGGCAGGAACATAACTGAGC | |
| D2176G-IDF | CAACCGTGAAGGATTACCTCTGT | Analysis of the D2176G mutation by Sanger sequencing |
| D2176G-IDR | GTGATTCTTCCCAGTCCACAAC | |
| W548L-F | GGAGTGAGTACGGTGTGCGCATTGAGAACCTCCCTGT | Analysis of the W548L mutation by NGS |
| W548L-R | GAGTTGGATGCTGGATGGATCTCGCTCTCACATTCCG | |
| S627I-F | GGAGTGAGTACGGTGTGCGCCGCCATCAAGAAGA | Analysis of the S627I mutation by NGS |
| S627I-R | GAGTTGGATGCTGGATGGCAGTCCTGCCATCACCAT | |
| I1879V-F | GGAGTGAGTACGGTGTGCAGTGGTGAAATTAGGTGGG | Analysis of the I1879V mutation by NGS |
| I1879V-R | GAGTTGGATGCTGGATGGCCAACAGTTCTTCCAGTCAC | |
| D2176G-F | GGAGTGAGTACGGTGTGCCCATGGCTGCAGAGCTACGA | Analysis of the D2176G mutation by NGS |
| D2176G-R | GAGTTGGATGCTGGATGGACCCTTGCGGTTCCAGAACA | |
| EPSPS-F | GGAGTGAGTACGGTGTGCGGTGGCAAGTTTCCTGTT | Analysis of the TAP mutation by NGS |
| EPSPS-R | GAGTTGGATGCTGGATGGCCCCATGAATTCCATACAT | |

### Supplemental sequence 1. MCP1-RT

**NLS-MCP1-linker-M_MLV_RT-NLS**

ATGCCTAAGAAGAAGAGGAAGGTGGGCTCAATGGCGTCCAACTTCACCCAGTTCGTGCTGGTCGACAACGGCGGCACGGGCGACGTGACCGTCGCT
CCAAGCAACTTCGCTAACGGCATCGCCGAGTGGATCTCCAGCAACTCCAGGAGTCAGGCGTACAAGGTGACGTGCTCAGTCAGGCAGTCCAGCGCTC
AGAACCGCAAGTACACCATCAAGGTCGAGGTCCCGAAGGGTGCGTGGCGCTCATACCTCAACATGGAGCTGACGATCCCGATCTTCGCGACCAACAG
CGACTGCGAGCTGATCGTGAAGGCCATGCAGGGCCTCCTGAAGGACGGCAACCCAATCCCAAGCGCTATCGCTGCGAACTCCGGCATCTACTCAGGC
GGCTCATCGGGCGGGTCAAGCGGGTCGGAGACACCGGGCACATCAGAGAGCGCTACCCCTGAGTCATCAGGCGGCTCTTCAGGCGGCAGCTCAACC
CTGAACATTGAGGACGAGTACCGGCTGCACGAGACGAGCAAGGAGCCAGACGTTTCGCTCGGCAGCACTTGGCTCTCTGACTTCCCACAGGCTTGG
GCCGAGACTGGCGGCATGGGCCTGGCCGTGCGCCAGGCTCCACTGATCATCCCTCTGAAGGCGACCTCCACCCCGGTTTCTATTAAGCAGTACCCGAT
GAGCCAGGAGGCCAGGCTGGGGATCAAGCCACACATTCAGCGGCTGCTGGACCAGGGCATCCTGGTGCCATGCCAGTCCCCGTGGAATACTCCGCTC
CTGCCGGTGAAGAAGCCTGGGACAAACGACTACAGGCCGGTTCAGGATCTCAGGGAGGTGAACAAGCGCGTGGAGGACATCCATCCGACAGTGCC
GAACCCGTACAATCTGCTGTCGGGCCTGCCTCCGAGCCACCAGTGGTACACCGTCCTGGACCTCAAGGACGCTTTCTTCTGCCTGCGGCTGCACCCGA
CGTCTCAGCCGCTGTTCGCGTTCGAGTGGCGCGACCCAGAGATGGGCATTTCCGGCCAGCTGACCTGGACACGCCTACCCCAGGGCTTCAAGAACTC
CCCGACTCTCTTCAACGAGGCTCTCCACCGGGATCTCGCGGACTTCAGGATTCAGCATCCCGATCTGATCCTGCTCCAGTATGTTGACGACCTCCTCCTG
GCCGCGACGTCGGAGCTGGACTGCCAGCAGGGCACCCGGGCGCTGCTGCAGACACTGGGCAATCTGGGGTACCGCGCCTCTGCGAAGAAGGCGCA
GATCTGCCAGAAGCAAGTGAAGTACCTGGGCTACCTCCTGAAGGAGGGCCAGCGCTGGCTCACTGAGGCGAGGAAGGAGACTGTTATGGGCCAGCC
CACTCCAAAGACTCCGAGGCAGCTCAGGGAGTTCCTCGGCAAGGCTGGGTTCTGCCGCCTGTTCATCCCTGGGTTCGCTGAGATGGCTGCGCCGCTC
TACCCGCTGACTAAGCCGGGGACACTGTTCAACTGGGGGCCAGACCAGCAGAAGGCGTACCAGGAGATTAAGCAGGCGCTGCTGACGGCCCCAGCG
CTCGGCCTACCAGACCTGACGAAGCCGTTCGAGCTGTTCGTTGACGAGAAGCAGGGGTACGCGAAGGGCGTGCTGACACAGAAGCTGGGGCCTTG
GCGCCGCCCGGTCGCGTACCTGTCGAAGAAGCTGGACCCAGTCGCTGCTGGGTGGCCTCCATGCCTCCGGATGGTCGCTGCTATTGCGGTTCTGACCA
AGGATGCGGGGAAGCTCACAATGGGGCAGCCTCTCGTGATCCTGGCTCCACATGCGGTGGAGGCGCTGGTGAAGCAGCCACCGGACCGGTGGCTGT
CGAACGCTCGGATGACACACTACCAGGCGCTCCTCCTCGATACAGACCGGGTTCAGTTCGGGCCTGTGGTTGCTCTGAACCCAGCCACACTGCTGCCA
CTCCCTGAGGAGGGCCTCCAGCACAATTGCCTCGACATCCTGGCTGAGGCGCACGGCACCCGCCCTGATCTCACCGACCAGCCTCTGCCAGATGCTGA
CCACACCTGGTACACGGATGGGTCCTCGCTGCTGCAGGAGGGCCAGAGGAAGGCGGGCGCCGCCGTCACCACAGAGACAGAGGTTATTTGGGCCA
AGGCCCTACCGGCTGGCACCAGCGCCCAGCGCGCTGAGCTGATCGCGCTGACTCAGGCGCTGAAGATGGCCGAGGGGAAGAAGCTCAATGTTTACA
CCGACTCGCGGTACGCGTTCGCTACAGCTCACATTCATGGGGAGATCTACCGCCGGCGCGGGTGGCTGACTTCGGAGGGCAAGGAGATTAAGAATAA
GGACGAGATCCTGGCCCTGCTCAAGGCGCTGTTCCTGCCGAAGCGCCTCTCAATCATTCACTGCCCGGGCCACCAGAAGGGCCATTCGGCCGAGGCT
AGGGGCAATCGGATGGCTGACCAGGCGGCGCGGAAGGCGGCTATCACCGAGACTCCCGATACATCTACCCTCCTGATCGAGAACTCGAGCCCAAGCG
GCGGGAGCAAGCGGACTGCGGATGGGTCTGAGTTCGAGCCAAAGAAGAAGAGGAAGGTGTGA

The codons in yellow font in MCP1 are different from those in MCP2.

### Supplemental sequence 2. MCP2-RT

**MCP2-linker-M_MLV_RT-NLS**

ATGGCGTCCAACTTCACCCAGTTCGTGCTGGTCGACAACGGCGGCACGGGCGACGTGACCGTCGCTCCAAGCAACTTCGCTAACGGCGTGGCCGAGT
GGATCTCCAGCAACTCCAGGAGTCAGGCGTACAAGGTGACGTGCTCAGTCAGGCAGTCCAGCGCTCAGAAGCGCAAGTACACCATCAAGGTCGAGG
TCCCGAAGGTGGCCACGCAGACCGTGGGCGGCGTCGAGCTGCCAGTCGCTGCGTGGCGCTCATACCTCAACATGGAGCTGACGATCCCGATCTTCGC
GACCAACAGCGACTGCGAGCTGATCGTGAAGGCCATGCAGGGCCTCCTGAAGGACGGCAACCCAATCCCAAGCGCTATCGCTGCGAACTCCGGCATC
TACTCAGGCGGCTCATCGGGCGGGTCAAGCGGGTCGGAGACACCGGGCACATCAGAGAGCGCTACCCCTGAGTCATCAGGCGGCTCTTCAGGCGGC
AGCTCAACCCTGAACATTGAGGACGAGTACCGGCTGCACGAGACGAGCAAGGAGCCAGACGTTTCG//AAGGCGGCTATCACCGAGACTCCCGATAC
ATCTACCCTCCTGATCGAGAACTCGAGCCCAAGCGGCGGGAGCAAGCGGACTGCGGATGGGTCTGAGTTCGAGCCAAAGAAGAAGAGGAAGGTGT
GA

The codons in yellow font in MCP2 are different from those in MCP1. “//” indicates omitted sequence which is the same as the above.

### Supplemental sequence 3. MCP-RT-MCP

**NLS-MCP1-linker-M_MLV_RT-linker-MCP1b-NLS**

ATGCCTAAGAAGAAGAGGAAGGTGGGCTCAATGGCGTCCAACTTCACCCAGTTCGTGCTGGTCGACAACGGCGGCACGGGCGACGTGACCGTCGCT
CCAAGCAACTTCGCTAACGGCATCGCCGAGTGGATCTCCAGCAACTCCAGGAGTCAGGCGTACAAGGTGACGTGCTCAGTCAGGCAGTCCAGCGCTC
AGAACCGCAAGTACACCATCAAGGTCGAGGTCCCGAAGGGTGCGTGGCGCTCATACCTCAACATGGAGCTGACGATCCCGATCTTCGCGACCAACAG
CGACTGCGAGCTGATCGTGAAGGCCATGCAGGGCCTCCTGAAGGACGGCAACCCAATCCCAAGCGCTATCGCTGCGAACTCCGGCATCTACTCAGGC
GGCTCATCGGGCGGGTCAAGCGGGTCGGAGACACCGGGCACATCAGAGAGCGCTACCCCTGAGTCATCAGGCGGCTCTTCAGGCGGCAGCTCAACC
CTGAACATTGAGGACGAGTACCGGCTGCACGAGACGAGCAAGGAGCCAGACGTTTCG//AAGGCGGCTATCACCGAGACTCCCGATACATCTACCCTC
CTGATCGAGAACTCGAGCCCATCAGGCGGGTCGTCAGGCGGCAGCTCAGGCTCAGAGACACCTGGCACCAGCGAGTCAGCTACACCGGAGTCATCT
GGCGGGTCATCTGGCGGGTCGTCAATGGCCAGCAACTTCACGCAATTTGTGCTCGTGGACAACGGCGGCACGGGCGACGTAACCGTGGCTCCGTCCA
ACTTCGCGAACGGGATCGCCGAGTGGATCAGCAGCAACTCCCGCAGCCAGGCGTACAAGGTGACCTGCAGCGTCAGGCAGTCCAGCGCACAGAATC
GCAAGTACACGATCAAGGTGGAGGTACCGAAGGGGGCCTGGCGCTCATACCTCAATATGGAGCTGACTATTCCTATTTTCGCCACGAACTCCGATTGC

GAGCTGATCGTGAAGGCGATGCAGGGCCTCCTCAAGGATGGGAACCCGATCCCATCGGCGATCGCTGCGAACAGCGGGATCTACAGCGGCGGGAGC
AAGCGGACTGCGGATGGGTCTGAGTTCGAGCCAAAGAAGAAGAGGAAGGTGTGA
"//" indicates omitted sequence which is the same as the above.

### Supplemental sequence 4. RT-MCP

**NLS-M_MLV_RT-linker-MCP1b-NLS**

ATGAAGAGGACAGCCGATGGCAGCGAGTTCGAGAGCCCTAAGAAGAAGAGGAAGGTGACCCTGAACATTGAGGACGAGTACCGGCTGCACGAGAC
GAGCAAGGAGCCAGACGTTTCG//AAGGCGGCTATCACCGAGACTCCCGATACATCTACCCTCCTGATCGAGAACTCGAGCCCATCAGGCGGGTCGTC
AGGCGGCAGCTCAGGCTCAGAGACACCTGGCACCAGCGAGTCAGCTACACCGGAGTCATCTGGCGGGTCATCTGGCGGGTCGTCAATGGCCAGCAA
CTTCACGCAATTTGTGCTCGTGGACAACGGCGGCACGGGCGACGTAACCGTGGCTCCGTCCAACTTCGCGAACGGGATCGCCGAGTGGATCAGCAGC
AACTCCCGCAGCCAGGCGTACAAGGTGACCTGCAGCGTCAGGCAGTCCAGCGCACAGAATCGCAAGTACACGATCAAGGTGGAGGTACCGAAGGGG
GCCTGGCGCTCATACCTCAATATGGAGCTGACTATTCCTATTTTCGCCACGAACTCCGATTGCGAGCTGATCGTGAAGGCGATGCAGGGCCTCCTCAAG
GATGGGAACCCGATCCCATCGGCGATCGCTGCGAACAGCGGGATCTACAGCGGCGGGAGCAAGCGGACTGCGGATGGGTCTGAGTTCGAGCCAAAG
AAGAAGAGGAAGGTGTGA
"//" indicates omitted sequence which is the same as the above.

### Supplemental sequence 5. SpCas9H840A-P2A-MR

**zCas9H840A-P2A- NLS-MCP1-linker-M_MLV_RT-NLS**

//CTCGGGGGCGACAAGCGGCCAGCGGCGACGAAGAAGGCGGGGCAGGCGAAGAAGAAGAAGGGAAGCGGAGCTACTAACTTCAGCCTGCTGAA
GCAGGCTGGAGACGTGGAGGAGAACCCTGGACCTATGCCTAAGAAGAAGAGGAAGGTGGGCTCAATGGCGTCCAACTTCACCCAGTTCGTGCTGGT
CGACAACGGCGGCACGGGCGACGTGACC//
"//" indicates omitted sequence which is the same as the above.

### Supplemental sequence 6. MCP-CaMV_RT

**NLS-MCP1-linker-CaMV_RT-NLS**

ATGCCTAAGAAGAAGAGGAAGGTGGGCTCAATGGCGTCCAACTTCACCCAGTTCGTGCTGGTCGACAACGGCGGCACGGGCGACGTGACCGTCGCT
CCAAGCAACTTCGCTAACGGCATCGCCGAGTGGATCTCCAGCAACTCCAGGAGTCAGGCGTACAAGGTGACGTGCTCAGTCAGGCAGTCCAGCGCTC
AGAACCGCAAGTACACCATCAAGGTCGAGGTCCCGAAGGGTGCGTGGCGCTCATACCTCAACATGGAGCTGACGATCCCGATCTTCGCGACCAACAG
CGACTGCGAGCTGATCGTGAAGGCCATGCAGGGCCTCCTGAAGGACGGCAACCCAATCCCAAGCGCTATCGCTGCGAACTCCGGCATCTACTCAGGC
GGCTCATCGGGCGGGTCAAGCGGGTCGGAGACACCGGGCACATCAGAGAGCGCTACCCCTGAGTCATCAGGCGGCTCTTCAGGCGGCAGCTCCATG
GATCATCTGCTCCTCAAGACACAGACCCAGACCGAGCAAGTGATGAATGTGACCAACCCGAACTCCATCTACATCAAGGGGCGCCTCTATTTCAAGGG
CTATAAGAAGATCGAACTCCATTGCTTTGTGGACACCGGCGCGTCCCTCTGCATCGCGAGCAAGTTCGTCATCCCAGAAGAGCATTGGGTGAACGCGG
AGAGGCCGATCATGGTGAAGATCGCCGACGGCTCCTCCATCACCATCAGCAAGGTGTGCAAGGATATCGATCTCATCATCGCCGGCGAGATCTTCCGCA
TTCCAACCGTCTACCAGCAAGAGTCCGGCATCGACTTTATTATCGGCAACAACTTCTGCCAGCTGTACGAACCGTTCATCCAGTTCACCGACCGCGTGAT
CTTCACAAAGAACAAGTCCTACCCGGTCCACATCGCCAAACTGACAAGGGCGGTGAGGGTGGGGACAGAGGGCTTTCTCGAGAGCATGAAGAAGA
GGAGCAAGACACAGCAGCCAGAGCCGGTGAATATCAGCACCAACAAGATCGAAAACCCGCTGGAGGAGATTGCCATTCTCAGCGAGGGGAGGAGG
CTCTCCGAGGAGAAGCTCTTTATTACCCAGCAGCGCATGCAGAAGATCGAAGAACTGCTCGAGAAAGTCTGCTCCGAGAACCCACTCGACCCGAATAA
GACCAAGCAGTGGATGAAGGCGAGCATCAAACTCTCCGATCCGAGCAAAGCGATCAAAGTGAAGCCGATGAAGTACTCCCCGATGGATCGCGAGGA
GTTCGACAAGCAGATTAAGGAGCTGCTGGACCTCAAAGTCATCAAGCCGTCCAAATCCCCGCACATGGCCCCAGCCTTTCTCGTGAACAACGAGGCG
GAGAAGAGGAGGGGCAAGAAGAGGATGGTCGTGAACTACAAGGCGATGAACAAAGCGACCGTGGGGGATGCGTATAATCTGCCGAACAAGGACGA
GCTGCTCACACTGATTCGCGGCAAGAAGATCTTCTCCAGCTTCGATTGCAAGTCAGGGTTCTGGCAAGTGCTCCTCGACCAAGAATCTCGCCCACTCAC
CGCCTTTACATGCCCGCAAGGCCACTACGAGTGGAACGTCGTGCCATTCGGGCTCAAGCAAGCCCCAAGCATCTTCCAGCGCCACATGGATGAAGCCT
TCCGCGTGTTCCGCAAATTCTGCTGCGTCTACGTCGATGACATCCTCGTGTTTTCCAACAACGAGGAGGACCACCTCCTCCATGTGGCCATGATTCTCCA
GAAGTGCAATCAGCACGGCATCATTCTCTCAAAGAAGAAAGCGCAGCTGTTCAAGAAGAAGATCAACTTTCTCGGCCTCGAGATTGATGAGGGCACC
CATAAGCCGCAAGGCCATATTCTCGAGCACATCAACAAGTTTCCGGACACCCTCGAGGACAAGAAGCAACTGCAACGCTTTCTCGGCATCCTCACATAC
GCGAGCGACTATATCCCAAAACTCGCCCAGATCCGCAAGCCACTCCAAGCCAAACTGAAGGAAAACGTCCCGTGGCGCTGGACCAAAGAGGACACCC
TCTACATGCAGAAAGTGAAGAAAAATCTCCAAGGCTTTCCGCCGCTGCACCATCCACTCCCGGAGGAAAAGCTCATCATTGAAACCGACGCGAGCGAT
GATTACTGGGGCGGGATGCTCAAGGCCATCAAGATTAACGAGGGCACCAACACCGAACTCATCTGTCGCTACGCGTCAGGGTCCTTTAAGGCGGCCG
AGAAGAACTACCACAGCAACGATAAGGAAACCCTCGCGGTCATCAACACCATCAAAAAGTTCAGCATCTATCTCACCCCGGTGCACTTTCTCATCCGCA
CCGACAACACACACTTTAAGAGCTTCGTCAACCTCAACTATAAGGGCGATTCCAAGCTCGGGCGCAACATTAGGTGGCAAGCGTGGCTCTCACACTAC
AGCTTTGACGTCGAGCACATCAAGGGCACCGACAACCATTTCGCCGACTTTCTCAGCCGCGAGTTCAACAAGGTCAATTCCAGCGGCGGGAGCAAGC
GGACTGCGGATGGGTCTGAGTTCGAGCCAAAGAAGAAGAGGAAGGTGTGA

#### Supplemental sequence 7. MCP-SS4_RT

**NLS-MCP1-linker-SS4_RT-NLS**

ATGCCTAAGAAGAAGAGGAAGGTGGGCTCAATGGCGTCCAACTTCACCCAGTTCGTGCTGGTCGACAACGGCGGCACGGGCGACGTGACCGTCGCT
CCAAGCAACTTCGCTAACGGCATCGCCGAGTGGATCTCCAGCAACTCCAGGAGTCAGGCGTACAAGGTGACGTGCTCAGTCAGGCAGTCCAGCGCTC
AGAACCGCAAGTACACCATCAAGGTCGAGGTCCCGAAGGGTGCGTGGCGCTCATACCTCAACATGGAGCTGACGATCCCGATCTTCGCGACCAACAG
CGACTGCGAGCTGATCGTGAAGGCCATGCAGGGCCTCCTGAAGGACGGCAACCCAATCCCAAGCGCTATCGCTGCGAACTCCGGCATCTACTCAGGC
GGCTCATCGGGCGGGTCAAGCGGGTCGGAGACACCGGGCACATCAGAGAGCGCTACCCCTGAGTCATCAGGCGGCTCTTCAGGCGGCAGCTCAACC
CTGAACATTGAGGACGAGTACCGGCTGCACGAGACGAGCAAGGAGCCAGACGTTTCGCTCGGCAGCACTTGGCTCTCTGACTTCCCACAGGCTTGG
GCCGAGACTGGCGGCATGGGCCTGGCCGTGCGCCAGGCTCCACTGATCATCCTCCTGAAGGCGACCTCCACCCCGGTTTCTATTAAGCAGTACCCGAT
GCGGCAGAAGGCCAGGCTGGGGATCAAGCCACACATTCAGCGGCTGCTGGACCAGGGCATCCTGGTGCCATGCCAGTCCCCGTGGAATACTCCGCTC
CTGCCGGTGAAGAAGCCTGGGACAAACGACTACAGGCCGGTTCAGGATCTCAGGGAGGTGAACAAGCGCGTGGAGGACATCCATCCGACAGTGCC
GAACCCGTACAATCTGCTGTCGGGCCTGCCTCCGAGCCACCAGTGGTACACCGTCCTGGACCTCAAGGACGCTTTCTTCTGCCTGCGGCTGCACCCGA
CGTCTCAGCCGCTGTTCGCGTTCGAGTGGCGCGACCCAGAGATGGGCATTTCCGGCCAGCTGACCTGGACACGCCTACCCCAGGGCTTCAAGAACTC
CCCGGCGCTCTTCGACGAGGCTCTCCGCCGGGATCTCGCGGACTTCAGGATTCAGCATCCCGATCTGATCCTGCTCCAGTATGTTGACGACCTCCTCCT
GGCCGCGACGTCGGAGCTGGACTGCCAGCAGGGCACCCGGGCGCTGCTGCAGACACTGGGCGACCTGGGGTACCGCGCCTCTGCGAAGAAGGCGC
AGATCTGCCAGAAGCAAGTGAAGTACCTGGGCTACCTCCTGAAGGAGGGCCAGCGCTGGCTCACTGAGGCGAGGAAGGAGACTGTTATGGGCCAGC
CCACTCCAAAGACTCCGAGGCAGCTCAGGAAGTTCCTCGGCACTGCTGGGAACTGCCGCCTGTTCATCCCTGGGTTCGCTGAGATGGCTGCGCCGCT
CTACCCGCTGACTAAGCCGGGGACACTGTTCAACTGGGGGCCAGACCAGCAGAAGGCGTACCAGGAGATTAAGCAGGCGCTGCTGACGGCCCCAGC
GCTCGGCCTACCAGACCTGACGAAGCCGTTCGAGCTGTTCGTTGACGAGAAGCAGGGGTACGCGAAGGGCGTGCTGACACAGAAGCTGGGGCCTT
GGCGCCGCCCGGTCGCGTACCTGTCGAAGAAGCTGGACCCAGTCGCTGCTGGGTGGCCTCCATGCCTCCGGATGGTCGCTGCTATTGCGGTTCTGACC
AAGGATGCGGGGAAGCTCACAATGGGGCAGCCTCTCGTGATCGGCGCTCCACATGCGGTGGAGGCGCTGGTGAAGCAGCCACCGGACCGGTGGCT
GTCGAAGGCTCGGATGACACACTACCAGGCGCTCCTCCTCGATACAGACCGGGTTCAGTTCGGGCCTGTGGTTGCTCTGAACCCAGCCACACTGCTGC
CACTCCCTGAGGAGGGCCTCCAGCACAATTGCCTCGACATCCTGGCTGAGGCGCACGGCACCCGCCCTGATCTCACCGACCAGCCTCTGCCAGATGCT
GACCACACCTGGTACACGGGCGGGTCCTCGCTGCTGCAGGAGGGCCAGAGGAAGGCGGGCGCCGCCGTCACCACAGAGACAGAGGTTATTTGGGC
CAAGGCCCTACCGGCTGGCACCAGCGCCCAGCGCGCTGAGCTGATCGCGCTGACTCAGGCGCTGAGGATGGCCGAGGGGAAGAAGCTCAATGTTTA
CACCAACTCGCGGTACGCGTTCGCTACAGCTCACATTCAGGGGGAGATCTACCGCCGGCGCGGGCTCCTGACTTCGGAGGGCAAGGAGATTAAGAAT
AAGGACGAGATCCTGGCCCTGCTCAAGGCGCTGTTCCTGCCGAAGCGCCTCTCAATCATTCACTGCCCGGGCCACCAGAAGGGCCATTCGGCCGAGG
CTAGGGGCAATCGGATGGCTAACCAGGCGGCGCGGAAGGCGGCTATCACCGAGAACCCCGATACATCTACCCTCCCGATCGAGAACTCGAGCCCAAG
CGGCGGGAGCAAGCGGACTGCGGATGGGTCTGAGTTCGAGCCAAAGAAGAAGAGGAAGGTGTGA

SS4 RT harbors H8Y, P51L, S67R, E69K, T197A, H204R, N249D, E302K, F309N, W313F, T330P, L435G, N454K, D524G, K571R, D583N, H594Q, D653N, T664N, and L671P mutations (red letters) relative to wild-type M-MLV RT.

#### Supplemental sequence 8. Synthetic 2xMS2pegRs or 2xpegRNAs

**BsaI-MS2pegR/pegRNA-HDV-tMet-MS2pegR/pegRNA-BsaI**

**GGTCTCATGCANNNNNN//NNNNNNGGCCGGCATGGTCCCAGCCTCCTCGCTGGCGCCGGCTGGGCAACATGCTTCGGCATGGCGAATGGGAC**
**AACAACAAATCAGAGTGGCGCAGCGGAAGCGTGGTGGGCCCATAACCCACAGGTCCCAGGATCGAAACCTGGCTCTGATANNNNNN//NNNN**
**NNGGCCAGAGACC**

#### Supplemental sequence 9. Cassette expressing two MS2pegRs or pegRNAs

**35S-CmYLCV-U6-tGly- MS2pegR/pegRNA-HDV-tMet-MS2pegR/pegRNA-HDV-polyT-HSPt**

ATGGAGTCAAAGATTCAAATAGAGGACCTAACAGAACTCGCCGTAAAGACTGGCGAACAGTTCATACAGAGTCTCTTACGACTCAATGACAAGAAGAA
AATCTTCGTCAACATGGTGGAGCACGACACACTTGTCTACTCCAAAAATATCAAAGATACAGTCTCAGAAGACCAAAGGGCAATTGAGACTTTTCAACA
AAGGGTAATATCCGGAAACCTCCTCGGATTCCATTGCCCAGCTATCTGTCACTTTATTGTGAAGATAGTGGAAAAGGAAGGTGGCTCCTACAAATGCCA
TCATTGCGATAAAGGAAAGGCCATCGTTGAAGATGCCTCTGCCGACAGTGGTCCCAAAGATGGACCCCCACCCACGAGGAGCATCGTGGAAAAAGAA
GACGTTCCAACCACGTCTTCAAAGCAAGTGGATTGATGTGATTGGCAGACATACTGTCCCACAAATGAAGATGGAATCTGTAAAAGAAAACGCGTGAA
ATAATGCGTCTGACAAAGGTTAGGTCGGCTGCCTTTAATCAATACCAAAGTGGTCCCTACCACGATGGAAAAACTGTGCAGTCGGTTTGGCTTTTTCTG
ACGAACAAATAAGATTCGTGGCCGACAGGTGGGGGTCCACCATGTGAAGGCATCTTCAGACTCCAATAATGGAGCAATGACGTAAGGGCTTACGAAA
TAAGTAAGGGTAGTTTGGGAAATGTCCACTCACCCGTCAGTCTATAAATACTTAGCCCCTCCCTCATTGTTAAGGGAGCAAAATCTCAGAGAGATAGTCC
TAGAGAGAGAAAGAGAGCAAGTAGCCTAGAAGTAGTCAAGGCGGCGAAGTATTCAGGCACGTGGCCAGGAAGAAGAAAAGCCAAGACGACGAAA
ACAGGTAAGAGCTAAGCATCTAGATAAGTTGAAAACAATCTTCAAAAGTCCCACATCGCTTAGATAAGAAAACGAAGCTGAGTTTATATACAGCTAGAG
TCGAAGTAGTGATTGAACAAAGCACCAGTGGTCTAGTGGTAGAATAGTACCCTGCCACGGTACAGACCCGGGTTCGATTCCCGGCTGGTGCANNNNNN
N//NNNNNNGGCCGGCATGGTCCCAGCCTCCTCGCTGGCGCCGGCTGGGCAACATGCTTCGGCATGGCGAATGGGACAACAACAAATCAGAGTG
GCGCAGCGGAAGCGTGGTGGGCCCATAACCCACAGGTCCCAGGATCGAAACCTGGCTCTGATANNNNNN//NNNNNNGGCCGGCATGGTCCCAG
CCTCCTCGCTGGCGCCGGCTGGGCAACATGCTTCGGCATGGCGAATGGGACTTTTTTTTGATATCTCCGGGGCTAATTGAATATGAAGATGAAGATGAA
ATATTTGGTGTGTCAAATAAAAAGCTGGTGTGCTTAAGTTTGTGTTTTTTTCTTGGCTTGTTGTGTTATGAATTTGTGGCTTTTTCTAATATTAAATGAATG

TAAGATCTCATTATAATGAATAAACAAATGTTTCTATAATCCATTGTGAATGTTTTGTTGGATCTCTTCTGCAGCATATAACTACTGTATGTGCTATGGTATGG
ACTATGGAATATGATTAAAGATAAG

Note: the underlined part comes from a synthetic fragment.

#### Supplemental sequence 10. MS2pegR scaffolds

##### >pegRNA

NNNNNNNNNNNNNNNNNNNNGTTTAAGAGCTATGCTGGAAACAGCATAGCAAGTTTAAATAAGGCTAGTCCGTTATCAACTTGAAAAAGTGG
CACCGAGTCGGTGCNNNNNNNNNNNNNNNNNNNNNNNNNNNNN

##### >pegRNA2.6

NNNNNNNNNNNNNNNNNNNNGTTTAAGAGCTAGGCCAACATGAGGATCACCCATGTCTGCAGGGCCTAGCAAGTTTAAATAAGGCTAGTCCGTTA
TCAACTTGGCCAACATGAGGATCACCCATGTCTGCAGGGCCAAGTGGCACCGAGTCGGTGCNNNNNNNNNNNNNNNNNNNNNNNNNNNNN

##### >pegRNA2.3a

NNNNNNNNNNNNNNNNNNNNGTTTAAGAGCTATGCTGGAAACAGCATAGCAAGTTTAAATAAGGCTAGTCCGTTATCAACTTGAAAAAGTGGCAC
CGAGTCGGTGCGGGAGCACATGAGGATCACCCATGTGCCACGAGCGACATGAGGATCACCCATGTCGCTCGTGTTCCCNNNNNNNNNNNNNNNNN
NNNNNNNNNNNN

##### >pegRNA2f.3a

NNNNNNNNNNNNNNNNNNNNGTTTAAGAGCTATGCTGGAAACAGCATAGCAAGTTTAAATAAGGCTAGTCCGTTATCAACTTGAAAAAGTGGC
ACCGAGTCGGTGCGGGAGCACATGAGGATCACCCATGTGCGACTCCCACAGTCACTGGGGAGTCTTCCCNNNNNNNNNNNNNNNNNNNNNNNN
NNNNN

##### >pegRNA2.3b

NNNNNNNNNNNNNNNNNNNNGTTTAAGAGCTATGCTGGAAACAGCATAGCAAGTTTAAATAAGGCTAGTCCGTTATCAACTTGAAAAAGTGGC
ACCGAGTCGGTGCNNNNNNNNNNNNNNNNNNNNNNNNNNNNNGGGAGCACATGAGGATCACCCATGTGCCACGAGCGACATGAGGATCACCC
ATGTCGCTCGTGTTCCC

##### >pegRNA2f.3b

NNNNNNNNNNNNNNNNNNNNGTTTAAGAGCTATGCTGGAAACAGCATAGCAAGTTTAAATAAGGCTAGTCCGTTATCAACTTGAAAAAGTGGC
ACCGAGTCGGTGCNNNNNNNNNNNNNNNNNNNNNNNNNNNNNGGGAGCACATGAGGATCACCCATGTGCGACTCCCACAGTCACTGGGGAGT
CTTCCC

##### >pegRNA1.1

NNNNNNNNNNNNNNNNNNNNGTTTAAGAGCTAGGCCAACATGAGGATCACCCATGTCTGCAGGGCCTAGCAAGTTTAAATAAGGCTAGTCCGTTA
TCAACTTGAAAAAGTGGCACCGAGTCGGTGCNNNNNNNNNNNNNNNNNNNNNNNNNNNNN

##### >pegRNA1.2

NNNNNNNNNNNNNNNNNNNNGTTTAAGAGCTATGCTGGAAACAGCATAGCAAGTTTAAATAAGGCTAGTCCGTTATCAACTTGGCCAACATGA
GGATCACCCATGTCTGCAGGGCCAAGTGGCACCGAGTCGGTGCNNNNNNNNNNNNNNNNNNNNNNNNNNNNN

##### >pegRNA1.1.1

NNNNNNNNNNNNNNNNNNNNGTTTAAGAGCTATGCTGAACATGAGGATCACCCATGTCTGCAGCAGCATAGCAAGTTTAAATAAGGCTAGTCCGT
TATCAACTTGAAAAAGTGGCACCGAGTCGGTGCNNNNNNNNNNNNNNNNNNNNNNNNNNNNN

##### >pegRNA1.2.1

NNNNNNNNNNNNNNNNNNNNGTTTAAGAGCTATGCTGGAAACAGCATAGCAAGTTTAAATAAGGCTAGTCCGTTATCAACTTGCACGAGCGAC
ATGAGGATCACCCATGTCGCTCGTGCAAGTGGCACCGAGTCGGTGCNNNNNNNNNNNNNNNNNNNNNNNNNNNNN

##### >pegRNA1f.1

NNNNNNNNNNNNNNNNNNNNGTTTAAGAGCTATGCTGGACTCCCACAGTCACTGGGGAGTCCAGCATAGCAAGTTTAAATAAGGCTAGTCCGT
TATCAACTTGAAAAAGTGGCACCGAGTCGGTGCNNNNNNNNNNNNNNNNNNNNNNNNNNNNN

##### >pegRNA1f.2

NNNNNNNNNNNNNNNNNNNNGTTTAAGAGCTATGCTGGAAACAGCATAGCAAGTTTAAATAAGGCTAGTCCGTTATCAACTTGACTCCCACAGT
CACTGGGGAGTCAAGTGGCACCGAGTCGGTGCNNNNNNNNNNNNNNNNNNNNNNNNNNNNN

##### >pegRNA1.3a

NNNNNNNNNNNNNNNNNNNNGTTTAAGAGCTATGCTGGAAACAGCATAGCAAGTTTAAATAAGGCTAGTCCGTTATCAACTTGAAAAAGTGGC

ACCGAGTCGGTGCGCTCGAGCGACATGAGGATCACCCATGTCGCTCGAGCNNNNNNNNNNNNNNNNNNNNNNNNNNNNN

**>pegRNA1f.3a**

NNNNNNNNNNNNNNNNNNNNGTTTAAGAGCTATGCTGGAAACAGCATAGCAAGTTTAAATAAGGCTAGTCCGTTATCAACTTGAAAAAGTGGC
ACCGAGTCGGTGCGACTCCCACAGTCACTGGGGAGTCNNNNNNNNNNNNNNNNNNNNNNNNNNNNN

**>pegRNA1.3b**

NNNNNNNNNNNNNNNNNNNNGTTTAAGAGCTATGCTGGAAACAGCATAGCAAGTTTAAATAAGGCTAGTCCGTTATCAACTTGAAAAAGTGGC
ACCGAGTCGGTGCNNNNNNNNNNNNNNNNNNNNNNNNNNNNNCACGAGCGACATGAGGATCACCCATGTCGCTCGTG

**>pegRNA1f.3b**

NNNNNNNNNNNNNNNNNNNNGTTTAAGAGCTATGCTGGAAACAGCATAGCAAGTTTAAATAAGGCTAGTCCGTTATCAACTTGAAAAAGTGGC
ACCGAGTCGGTGCNNNNNNNNNNNNNNNNNNNNNNNNNNNNNGACTCCCACAGTCACTGGGGAGTC

**>pegRNA2f.6.1**

NNNNNNNNNNNNNNNNNNNNGTTTAAGAGCTATGCTGAACATGAGGATCACCCATGTCTGCAGCAGCATAGCAAGTTTAAATAAGGCTAGTCC
GTTATCAACTTGACTCCCACAGTCACTGGGGAGTCAAGTGGCACCGAGTCGGTGCNNNNNNNNNNNNNNNNNNNNNNNNNNNNN

**>pegRNA2f.6.2**

NNNNNNNNNNNNNNNNNNNNGTTTAAGAGCTATGCTGGACTCCCACAGTCACTGGGGAGTCCAGCATAGCAAGTTTAAATAAGGCTAGTCCGT
TATCAACTTGGCCAACATGAGGATCACCCATGTCTGCAGGGCCAAGTGGCACCGAGTCGGTGCNNNNNNNNNNNNNNNNNNNNNNNNNNNN
N

**>pegRNA3f.8.1**

NNNNNNNNNNNNNNNNNNNNGTTTAAGAGCTATGCTGGAAACAGCATAGCAAGTTTAAATAAGGCTAGTCCGTTATCAACTTGGCCAACATGA
GGATCACCCATGTCTGCAGGGCCAAGTGGCACCGAGTCGGTGCNNNNNNNNNNNNNNNNNNNNNNNNNNNNNGGGAGCACATGAGGATCA
CCCATGTGCGACTCCCACAGTCACTGGGGAGTCTTCCC

**>pegRNA3f.8.2**

NNNNNNNNNNNNNNNNNNNNGTTTAAGAGCTATGCTGGAAACAGCATAGCAAGTTTAAATAAGGCTAGTCCGTTATCAACTTGACTCCCACAGT
CACTGGGGAGTCAAGTGGCACCGAGTCGGTGCNNNNNNNNNNNNNNNNNNNNNNNNNNNNNGGGAGCACATGAGGATCACCCATGTGCGA
CTCCCACAGTCACTGGGGAGTCTTCCC

**>pegRNA4f.9.1**

NNNNNNNNNNNNNNNNNNNNGTTTAAGAGCTAGGCCAACATGAGGATCACCCATGTCTGCAGGGCCTAGCAAGTTTAAATAAGGCTAGTCCGT
TATCAACTTGGCCAACATGAGGATCACCCATGTCTGCAGGGCCAAGTGGCACCGAGTCGGTGCNNNNNNNNNNNNNNNNNNNNNNNNNNNN
NGGGAGCACATGAGGATCACCCATGTGCGACTCCCACAGTCACTGGGGAGTCTTCCC

**>pegRNA4f.9.2**

NNNNNNNNNNNNNNNNNNNNGTTTAAGAGCTATGCTGAACATGAGGATCACCCATGTCTGCAGCAGCATAGCAAGTTTAAATAAGGCTAGTCC
GTTATCAACTTGACTCCCACAGTCACTGGGGAGTCAAGTGGCACCGAGTCGGTGCNNNNNNNNNNNNNNNNNNNNNNNNNNNNNGGGAGCA
CATGAGGATCACCCATGTGCGACTCCCACAGTCACTGGGGAGTCTTCCC

**>pegRNA4f.9.3**

NNNNNNNNNNNNNNNNNNNNGTTTAAGAGCTATGCTGGACTCCCACAGTCACTGGGGAGTCCAGCATAGCAAGTTTAAATAAGGCTAGTCCGT
TATCAACTTGGCCAACATGAGGATCACCCATGTCTGCAGGGCCAAGTGGCACCGAGTCGGTGCNNNNNNNNNNNNNNNNNNNNNNNNNNNN
NGGGAGCACATGAGGATCACCCATGTGCGACTCCCACAGTCACTGGGGAGTCTTCCC

#### Supplemental sequence 11. Spacer, rtT, and PBS of MS2pegRs

**>OsACC-I1879V**

CAAGGAAGATGGACTTGGTGTGG (target)
ACTTCCATGTAcATTgTCgACAC/CAAGTCCATCTTC (rtT/PBS)

**>OsACC-D2176G**

ACGAGGAGGGGCTTGGGTTGTGG (target)
TTGCTAcCAACCACAA/CCCAAGCCCCT (rtT/PBS)

**>OsALS-W548L.a**

GGGTATGGTGGTGCAATGGGAGG (target)
AAACCTATCtTCta/ATTGCACCACCAT (rtT/PBS)

**>OsALS-W548L.b**

ATTTGGGTATGGTGGTGCAATGG (target)
ACCTATCCTCCaATTG/CACCACCATACCC (rtT/PBS)

**>OsALS-S627I.a**

CCTTGAATGCGCCCCCACTTGGG (target)
TGCCTATGATaCCAAt/TGGGGGCGCATTC (rtT/PBS)

**>OsALS-S627I.b**

GTGCTGCCTATGATCCCAAGTGG (target)
GAATGCGCCCCCAaTT/GGGATCATAGGCA (rtT/PBS)

**>OsEPSPS-TAP**

GCAGTCACGGCTGCTGTCAATGG (target)
TGGAAtTGtAATGCGAtCATTG/ACAGCAGCCGTGA (rtT/PBS)
